# Context-dependent requirement of H3K9 methyltransferase activity during cellular reprogramming to iPSCs

**DOI:** 10.1101/634949

**Authors:** Simon Vidal, Alexander Polyzos, Jorge Morales Valencia, Hongsu Wang, Emily Swanzey, Ly-sha Ee, Bhishma Amlani, Shengjiang Tu, Yixiao Gong, Valentina Snetkova, Jane A. Skok, Aristotelis Tsirigos, Sangyong Kim, Effie Apostolou, Matthias Stadtfeld

## Abstract

Methylation of histone 3 at lysine 9 (H3K9) is widely regarded as a major roadblock for cellular reprogramming and interference with associated methyltransferases such as EHMT1 and EHMT2 (also known as GLP and G9A, respectively) increases the efficiencies at which induced pluripotent stem cells (iPSCs) can be derived. Activation of histone and DNA demethylases by ascorbic acid (AA) has become a common approach to facilitate the extensive epigenetic remodeling required for iPSC formation, but possible functional interactions between the H3K9 methylation machinery and AA-stimulated enzymes remain insufficiently explored. Here we show that reduction of EHMT1/2 activity counteracts iPSC formation in an optimized reprogramming system in the presence of AA. Mechanistically, EHMT1/2 activity under these conditions is required for efficient downregulation of somatic genes and transition into an epithelial state. Of note, transient inhibition of EHMT1/2 during reprogramming yields iPSCs that fail to efficiently give rise to viable mice, suggesting persistent molecular defects in these cells. Genetic interference with the H3K9 demethylase KDM3B ameliorated the adverse effect of EHMT1/2 inhibition on iPSC formation. Together, our observations document novel functions of H3K9 methyltransferases during iPSC formation and suggest that the balancing of AA-stimulated enzymes by EHMT1/2 supports efficient and error-free iPSC reprogramming to pluripotency.

## INTRODUCTION

Covalent chromatin modifications such as DNA and histone methylation modulate gene expression and stabilize epigenetic states in a wide variety of biological processes. The genome-wide and locus-specific abundance of chromatin marks is determined by counteracting enzymatic activities, such as histone methyltransferases and demethylases (Black et al., 2012). Lysine methylation at position 9 of histone 3 (H3K9 methylation) is an epigenetic mark predominantly associated with gene repression that is conserved during evolution and plays important regulatory functions during embryonic development, sex determination, neuronal plasticity, immune cell function and tumorigenesis (Benevento et al., 2015; Casciello et al., 2015; Kuroki and Tachibana, 2018; Scheer and Zaph, 2017). H3K9 trimethylation in heterochromatic regions is established by SETDB1 and SUV39H1/2 (Kang, 2015; Peters et al., 2001; Rice et al., 2003), while the H3K9 mono- and dimethyltransferases EHMT1 and EHMT2 (also known as GLP and G9A, respectively) mediate gene silencing at euchromatic loci (Shinkai and Tachibana, 2011). Enzymes involved in the establishment of H3K9 methylation are repressors of core pluripotency-associated gene loci (Epsztejn-Litman et al., 2008) and, thus, have particular relevance for physiological and experimentally-induced changes in cell identity (Becker et al., 2016; Feldman et al., 2006). Inefficient removal of H3K9 methylation is a frequent cause of incomplete transcriptional reprogramming after somatic cell nuclear transfer (Matoba et al., 2014) and has been reported to impede the binding of reprogramming factors to the genome during the derivation of induced pluripotent stem cells (iPSCs) (Soufi et al., 2012). Consequently, the formation of mouse (Chen et al., 2013b; Liang et al., 2012; Sridharan et al., 2013; Tran et al., 2015; Wang et al., 2011; Wei et al., 2017) and human (Onder et al., 2012; Soufi et al., 2012) iPSCs can be substantially facilitated by interferences with H3K9 methyltransferases or by activation of respective demethylases. Despite the importance of H3K9 methyltransferases for cellular reprogramming and different physiological and pathological processes (Shankar et al., 2013), our understanding of the regulatory interactions controlling the function of these enzymes in different cellular contexts remains incomplete. We reasoned that the systematic comparison of cells that reprogram at markedly different efficiencies could be utilized to discover unexplored aspects of the H3K9 methylation machinery. By taking this approach we were able to assign specific and context-dependent functions to the histone methyltransferases EHMT1/2 during the reprogramming of fibroblasts into iPSCs by the “Yamanaka transcription factors (TFs)” OCT4, KLF4, SOX2 and MYC (OKSM). In particular, we report the unexpected finding that EHMT1/2 activity supports efficient and faithful establishment of pluripotency in conditions that favor histone demethylation.

## RESULTS

### Context-dependent roles of EHMT1/2 activity during mouse fibroblast reprogramming

We decided to compare the role of H3K9 methylation during iPSC formation driven solely by the OKSM factors (“basal reprogramming”) to OKSM-driven reprogramming supported by chemical modulation of the TGFβ and WNT signaling pathways (via iALK5 and CHIR99021, respectively) and of chromatin state (via ascorbic acid, AA) (“3c enhanced reprogramming”). These compounds are commonly used to facilitate reprogramming and pluripotent cell culture (Dakhore et al., 2018) and iPSCs generated in this manner are developmentally fully competent (Amlani et al., 2018). 3c reprogramming yields approximately 50 times more stable iPSC colonies than basal reprogramming and does so in a shorter period of time (six rather than twelve days of OKSM expression) (Penalosa-Ruiz et al., 2019; Saunders et al., 2017; Schwarz et al., 2018; Stelzer et al., 2015; Vidal et al., 2014). We therefore speculated that 3c might enhance iPSC formation by counteracting H3K9 methylation more efficiently than OKSM factors alone.

To our surprise, both H3K9 mono- and dimethylation (H3K9me1/2) levels increased shortly after initiation of OKSM expression (24 hours; 24h) in 3c conditions compared to uninduced mouse embryonic fibroblasts (MEFs), while a less pronounced increase was observed during basal reprogramming (**Figure 1A,B**). This coincided with increased protein levels of EHMT1 and EHMT2, the two major methyltransferases that catalyze H3K9me1/2 in mammalian cells (**Figure 1C** and **S1A**). We also observed transcriptional upregulation of all major H3K9 methyltransferases at this time while no such trend was observed with the corresponding demethylases (**Figure S1B**). The heterochromatic H3K9me3 mark (**Figure 1A,B**) as well as the activating marks H3K4me2 and H3K4me3 (**Figure S1C**) were reduced during basal and 3c reprogramming, supporting the notion that dynamic chromatin remodeling commences early during iPSC formation (Koche et al., 2011). EHMT1 and EHMT2 are two closely structurally related enzymes that work as heterodimers in pluripotent cells (Tachibana et al., 2005). Treatment of mouse embryonic fibroblasts (MEFs) undergoing reprogramming with standard doses (1 μM) of UNC0638, an efficient and specific substrate inhibitor of EHMT1/2 activity (Vedadi et al., 2011), greatly reduced levels of H3K9me2 (**Figure 1D**), demonstrating that the gain of this chromatin modification during early stages of iPSC formation is at least in part driven by active histone methylation. Together, these observations suggest that upregulation of components of the H3K9 methylation machinery is an early and previously unappreciated event during cellular reprogramming, which appears more pronounced in 3c enhanced compared to basal iPSC reprogramming conditions.

**Figure 1.**
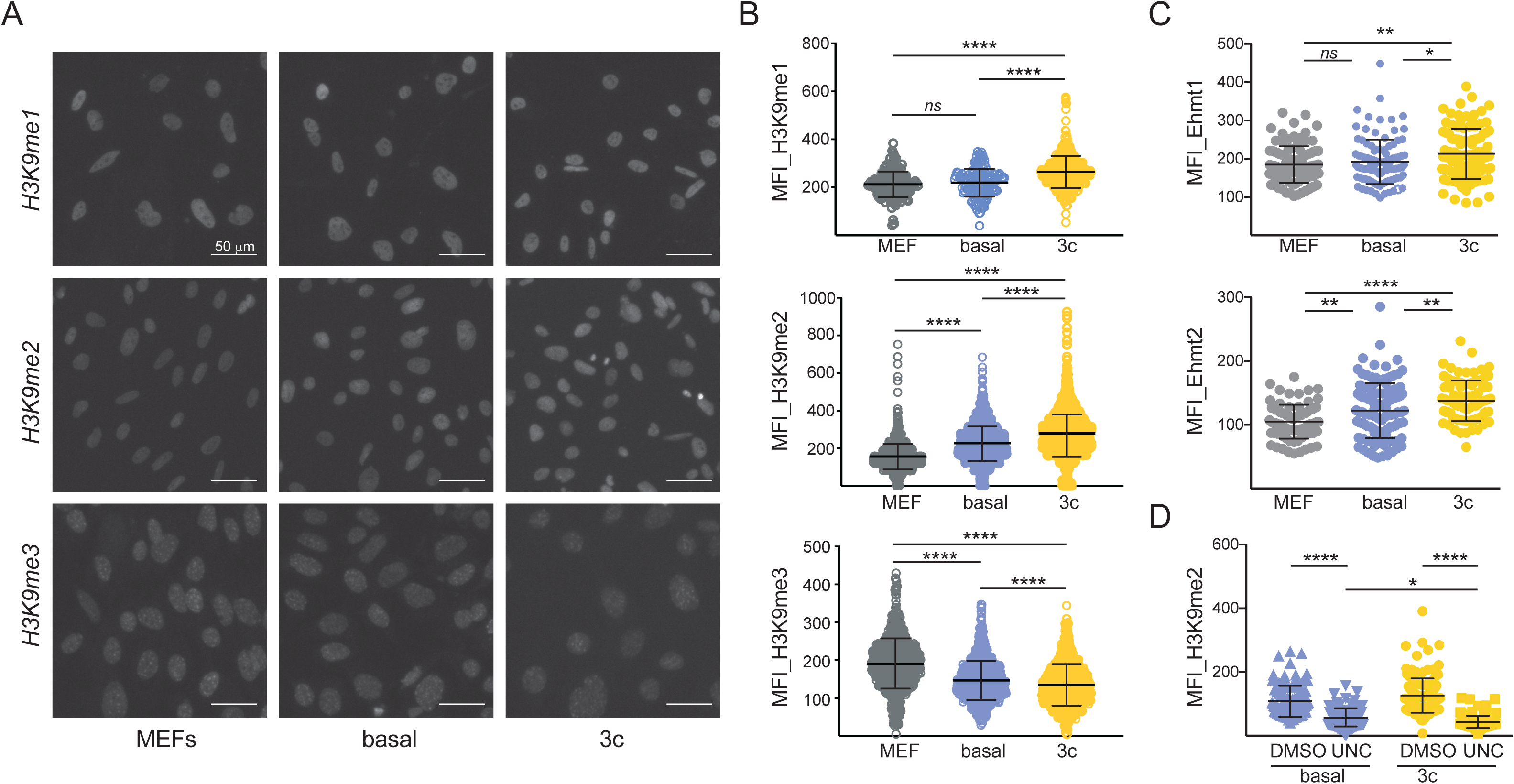
Increased EHMT1/2 activity during early stages of OKSM-driven reprogramming. **(A)** Fluorescence images of MEFs and cells 24h after initiation of reprogramming in indicated conditions after staining with antibodies against H3K9me1, H3K9me2 or H3K9me3, respectively. **(B)** Quantification of the mean fluorescence intensities of respective immunofluorescence stainings. More than 200 nuclei of similar size were quantified for each condition and antibody. **(C)** Same as (B) but after staining with antibodies against EHMT1 and EHMT2, respectively. Significance in (B-C) with one-way ANOVA with Tukey post-test with *p<0.05, **p<0.01 and ****p<0.0001**. (D)** Quantification of H3K9me2 levels in cells either exposed to UNC0638 (UNC) or DMSO for the first 24h of OKSM expression. Significance with one-way ANOVA with Sidak post-test *p<0.05 and ****p<0.0001

Chemical and genetic interference with EHMT1/2 during OKSM-driven reprogramming of mouse neural progenitor cells (Shi et al., 2008) and fibroblasts (Sridharan et al., 2013) has been reported to facilitate iPSC formation, supporting the prevalent notion that H3K9 methyltransferase activity counteracts the induction of pluripotency. Accordingly, EHMT1/2 inhibition via UNC0638 substantially increased the efficiency of iPSC formation from MEFs in basal conditions in our system (**Figure 2A,B**; upper panels). To our surprise, the formation of stable, transgene-independent iPSC colonies was strongly impaired (3-5 fold) when UNC0638 was administered during 3c enhanced reprogramming (**Figure 2A,B**; lower panels). We also observed significantly reduced colony numbers during 3c enhanced reprogramming when driving iPSC formation by OKS factors (no ectopic MYC) and when using constitutive lentiviral vectors to express OKSM (Sommer et al., 2009)(**Figure S2A-C**). Together, these observations suggest that, in contrast to basal reprogramming, 3c enhanced reprogramming does not counteract EHMT1/2, but instead partially becomes dependent on the activity of these enzymes.

**Figure 2.**
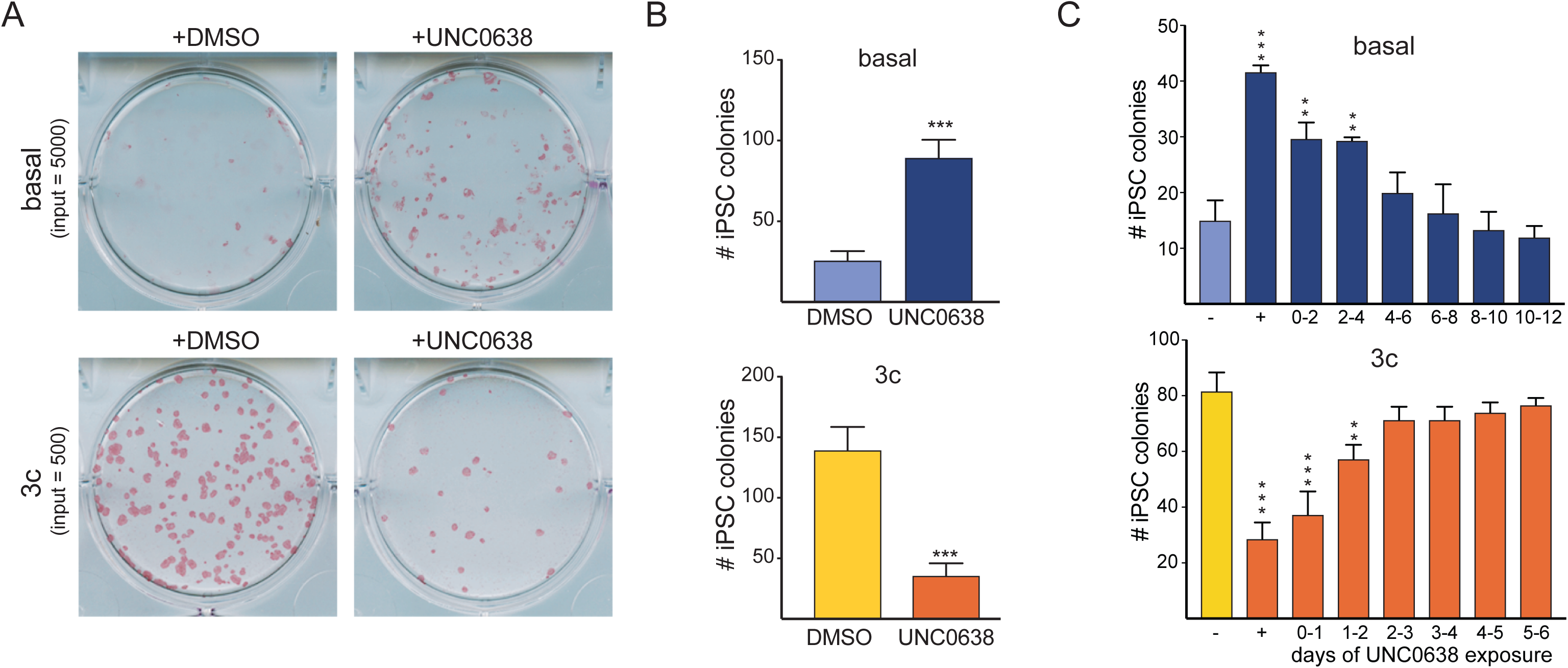
EHMT1/2 activity supports 3c enhanced reprogramming. **(A)** Alkaline phosphatase (AP) staining of iPSCs formed from indicated input MEFs via basal and 3c enhanced reprogramming in absence and presence of UNC0638, respectively. Images were taken four days after dox was removed to select for stably reprogrammed cells. **(B)** Quantification of iPSC colony formation in indicated conditions. n = 3 biological repeats. Significance with unpaired t-test with ***p<0.001. **(C)** Number of iPSC colonies formed from reprogramming cultures initiated in basal or 3c enhanced conditions that were exposed for indicated times to EHMT1/2 inhibitor. Input cells is 2500 (basal) and 300 (3c enhanced) MEFs, respectively. n = 3 biological repeats. Significance with one-way ANOVA with Dunnett post-test with **p<0.01 and ***p<0.001.

### Molecular and cellular changes modulated by EHMT1/2 activity during iPSC formation

Next, we exposed OKSM-expressing cells at specific intervals of time to compound inhibitor. This revealed that the reprogramming-promoting effect of EHMT1/2 inhibition on basal reprogramming as well as the reprogramming-counteracting effect of EHMT1/2 inhibition on 3c enhanced reprogramming was most pronounced when UNC0638 was administered during early reprogramming stages (days 0 to 4 during basal and days 0 to 2 during 3c enhanced reprogramming, respectively) (**Figure 2C**). To gain insight into the genome-wide transcriptional consequences of EHMT1/2 inhibition during the early stages of iPSC formation, we initially used RNA-sequencing (RNA-seq) to determine gene expression changes that distinguish basal from 3c enhanced reprogramming. K-means clustering of genes differentially expressed between starting MEFs and cells two days after initiation of OKSM expression under 3c conditions revealed four distinct groups of genes (**Figure 3A** and **Supplemental Table 1**), two of which distinguished between basal and 3c reprogramming (Clusters 2 and 3) while two others did not (Clusters 1 and 4). Specifically, silencing of genes associated with extracellular matrix remodeling and embryonic development (Cluster 1) and activation of genes driving DNA replication and chromatin organization (Cluster 4) were similarly efficient in basal and enhanced reprogramming. In contrast, strong activation of genes associated with RNA and energy metabolism (Cluster 2) and efficient silencing of genes required for cell adhesion (Cluster 4) were only observed in presence of 3c (**Figure 3A** and **Supplemental Table 2**). These observations document that basal and 3c enhanced reprogramming exhibit distinct molecular characteristics early during iPSC formation.

**Figure 3.**
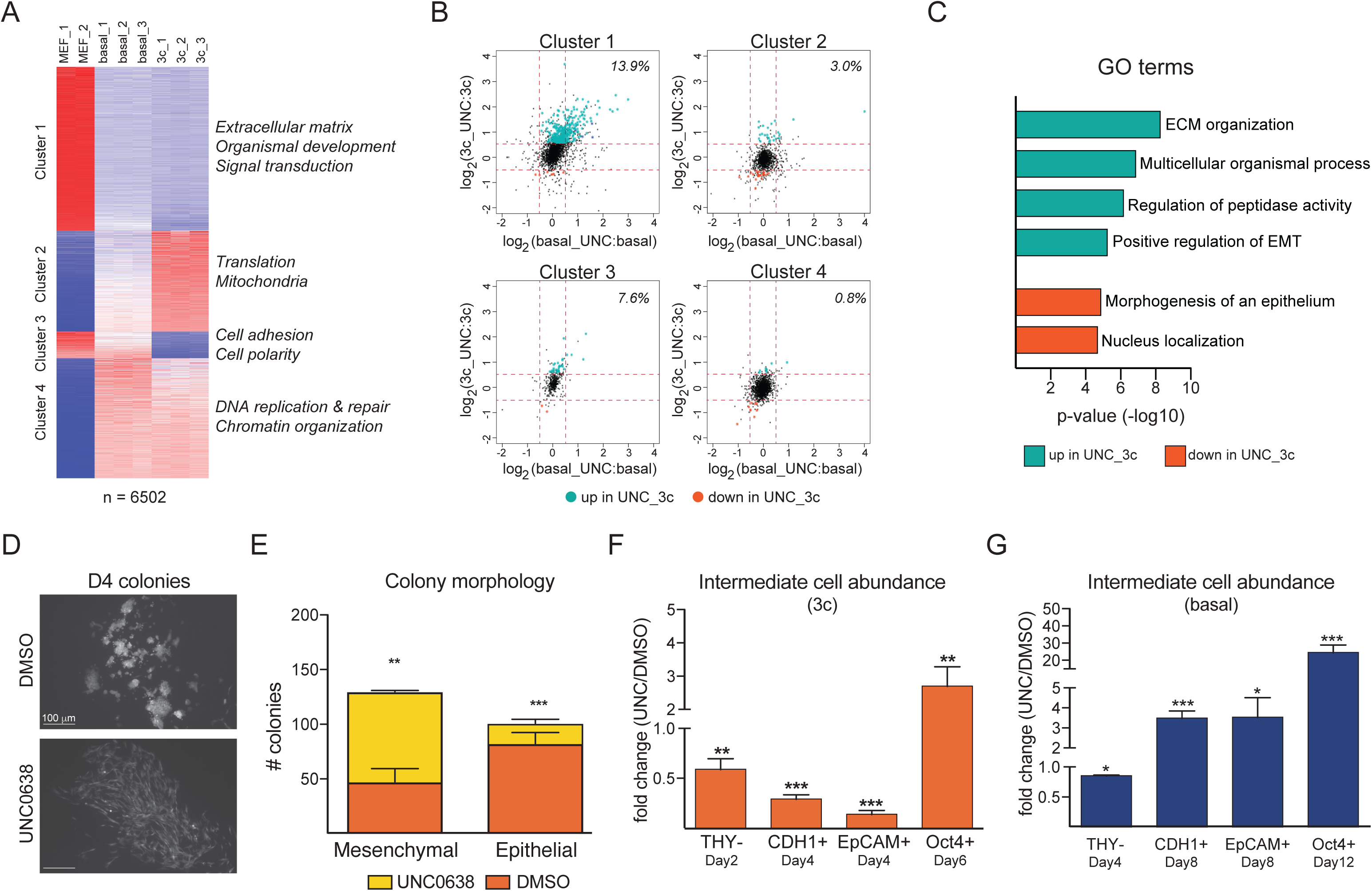
Molecular consequences of EHMT1/2 inhibition during early reprogramming stages. **(A)** K-means clustering of genes differentially expressed between MEFs and cells expressing OKSM for two days in 3c conditions (p-adj<0.05; FC>2). GO categories strongly associated with the four identified clusters of gene expression are highlighted. **(B)** Representation of the effect of EHMT1/2 inhibition (UNC) on the expression levels of genes associated with the four clusters identified in (A) during both basal (X axis) and 3c enhanced (Y axis) reprogramming. Transcripts with significant changed abundance during enhanced reprogramming (p-adj<0.05; FC>1.5) are highlighted in green (failed downregulation) and orange (failed upregulation), respectively. Numbers indicate percentage of cluster-specific genes affected by UNC0638 (UNC). **(C)** GO terms associated with gene loci that experience impaired activation (green) or impaired repression (orange) during 3c enhanced reprogramming in presence of EHMT1/2 inhibitor. **(D)** Representative images of colonies formed upon four days (D4) of OKSM expression in 3c conditions in absence (DMSO) or presence of UNC0638. Starting MEFs were labeled with ubiquitous EGFP to visualize colony morphology. **(E)** Quantification of D4 colonies with mesenchymal or epithelial morphology under indicated conditions. Significance with multiple two-tailed t-test with **p<0.01 and ***p<0.001. At least 100 colonies were scored in biological triplicate for each condition. **(F)** Relative abundance in presence versus absence of UNC0638 (UNC) of indicated reprogramming intermediates during 3c enhanced reprogramming, as measured by flow cytometry. **(G)** Same as (F) for reprogramming intermediates during basal reprogramming. Significance in (F,G) with multiple t-tests using Holm-Sidak correction with *p<0.05, **p<0.01 and ***p<0.001. N = 3 biological replicates.

Next, we interrogated the consequences of EHMT1/2 inhibition on the transcriptional dynamics associated with the two different reprogramming regimens. Principal component analysis (PCA) of early reprogramming intermediates confirmed close proximity of biological replicates and suggested a more pronounced effect of UNC0638 on enhanced than on basal reprogramming (**Figure S3A**). Differential gene expression analysis confirmed this observation and identified a total of 455 genes associated with efficient reprogramming that were significantly altered in their expression levels (fold change>1.5; p<0.05) in presence of UNC0638 (**Supplemental Table 3**). Almost none of these genes were affected at this stage of basal reprogramming. In agreement with the established roles of EHMT1/2 as transcriptional repressors we observed more frequent – but not exclusive – gene activation upon inhibition of these enzymes (**Figures 3B**). Further analysis of affected genes in the aforementioned four clusters revealed that EHMT1/2 inhibition counteracted the downregulation of a subset of fibroblastic genes (Cluster 1) (**Figure 3B** and **Supplemental Table 3**), suggesting that these enzymes contribute to the silencing of somatic gene expression during iPSC formation, similar to what was recently reported for other repressive complexes (Li et al., 2017). Gene ontology (GO) analysis supported this conclusion and suggested that EHMT1/2 inhibition interferes with cellular processes required for mesenchymal-to-epithelial transition (**Figure 3C** and **Supplemental Table 4**), an essential intermediate state of iPSC formation (Li et al., 2010; Samavarchi-Tehrani et al., 2010). Accordingly, we observed a pronounced reduction of nascent colonies with epithelial features (**Figure 3D,E**) and reduced numbers of intermediate cells expressing the epithelial surface markers CDH1 (also known as E-cadherin) and EpCAM (also known as CD326) during enhanced reprogramming when EHMT1/2 were inhibited (**Figure 3F**). In addition, the intermediate cells that successfully upregulated CDH1 retained elevated levels of the fibroblast surface marker THY1 (**Figure S3c**). In contrast, EHMT1/2 inhibition during basal reprogramming led to a significant increase of intermediate cells expressing CDH1 and EpCAM (**Figure 3G**). The relative abundance of cells that had reactivated expression of a reporter gene inserted into the endogenous *Pou5f1* locus (encoding *Oct4*) (Lengner et al., 2007) was moderately increased in 3c enhanced and dramatically increased in basal reprogramming (**Figure 3F,G**), which agrees with the established role of the H3K9 methylation machinery in silencing pluripotency-associated loci during development (Epsztejn-Litman et al., 2008; Feldman et al., 2006). These observations indicate that EHMT1/2 activity has distinct roles during basal and enhanced reprogramming. While UNC0638 treatment facilitates the reactivation of endogenous pluripotency loci in basal conditions, it counteracts the transition of reprogrammable fibroblast to intermediate, epithelialized stages of iPSC formation under optimized conditions. Of note, knockdown of *Ehmt1* recapitulated the effect of EHMT1/2 inhibition during enhanced reprogramming, while knockdown of *Ehmt2* recapitulated the effect of EHMT1/2 inhibition during basal reprogramming (**Figure S3D-F**), suggesting separate functions of these enzymes.

### EHMT1/2 activity balances AA-stimulated H3K9 demethylases during reprogramming

Next, we sought to investigate the molecular mechanisms responsible for the reduction of iPSC formation observed upon EHMT1/2 inhibition. Reprogramming experiments conducted in presence of single compounds revealed a significant reduction in iPSC colonies when UNC0638 was used together with AA (**Figure 4A,B**). In contrast, a slight increase in colony numbers was observed when EHMT1/2 were inhibited in presence of either CHIR99021 or iALK5 (**Figure 4B**). This reduction in colony number observed in presence of AA alone was less dramatic than in context of 3c, raising the possibility that AA might synergize with one or both of the other two reprogramming enhancing compounds in establishing a requirement for EHMT1/2 activity. Indeed, we observed a more pronounced reduction in iPSC colony numbers upon EHMT1/2 interference when AA was combined with iALK5 than with CHIR99021, while colony numbers remained unchanged when iALK5 was combined with CHIR99021 in absence of AA (**Figure S4A**). EHMT1/2 inhibition has been reported to counteract cellular proliferation in cancer cells (Casciello et al., 2015). In light of the importance of proliferative potential for iPSC formation (Li et al., 2009a; Utikal et al., 2009), we asked whether this could explain the observed reduction in colonies. Quantification of cells at early reprogramming stages revealed reduced cell numbers in presence of UNC0638 in all conditions, including those where EHMT1/2 inhibition favors reprogramming (**Figure S4B**). This suggests that an effect on proliferation cannot explain the reduction in iPSC reprogramming observed only in specific conditions.

**Figure 4.**
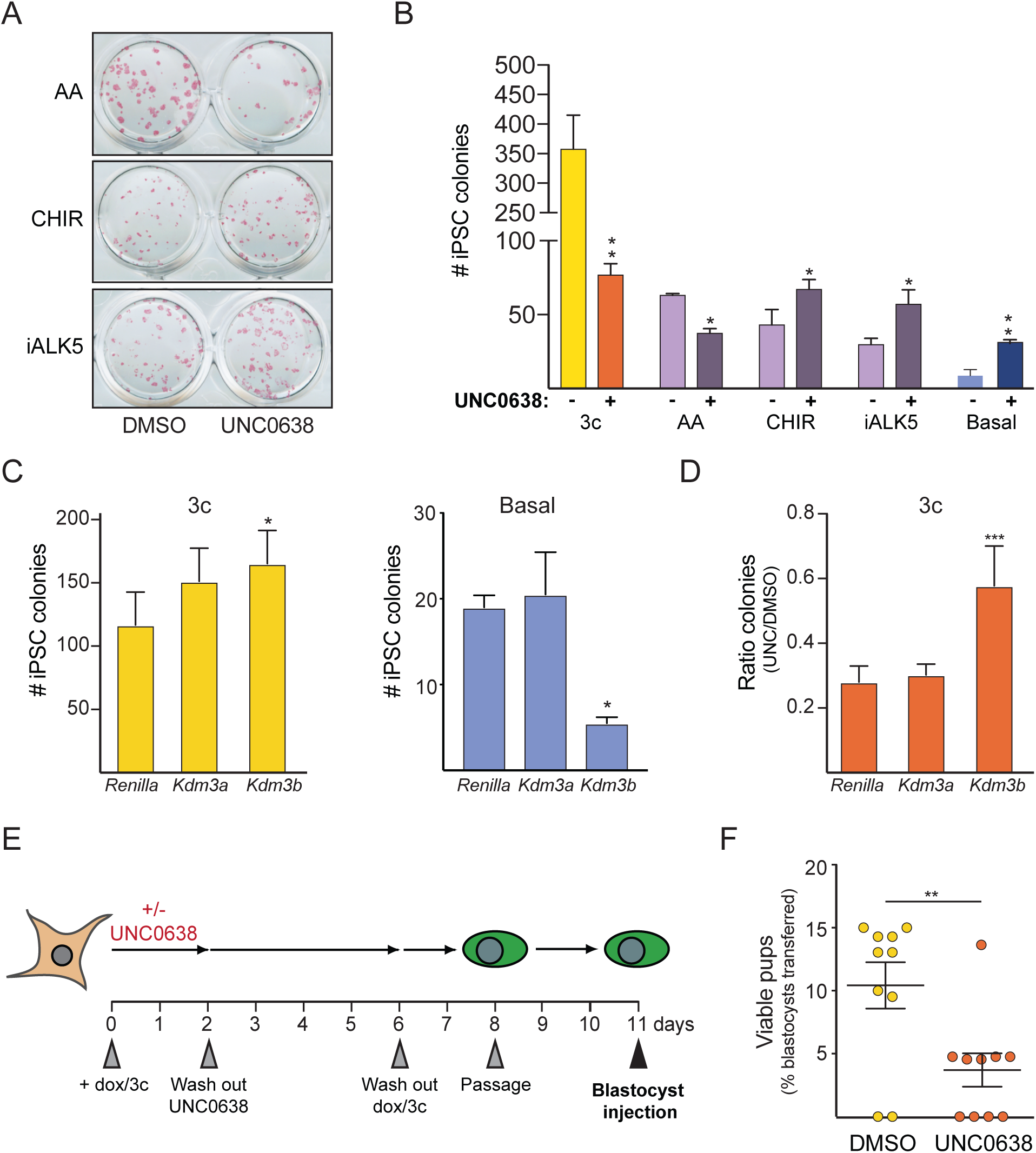
AA establishes a requirement for EHMT1/2 activity during enhanced iPSC reprogramming. **(A)** AP staining of transgene-independent iPSC colonies obtained after reprogramming MEFs in presence of indicated compounds and in absence or presence of UNC0638, respectively. **B)** Quantification of iPSC colonies formed in indicated conditions. N = 3 biological repeats. Significance was calculated by multiple t-test with p-values of *<0.05 and **<0.01. **(C)** Number of iPSC colonies formed after shRNA-mediated KD of *Kdm3a* or *Kdm3b* during 3c enhanced (left panel) and basal reprogramming (right panel), respectively. **(D)** Ratio of iPSC colonies formed during 3c enhanced reprogramming in presence and absence of EHMT1/2 inhibitor upon KD of indicated H3K9 demethylases. Significance in (C-E) with one-way ANOVA with Dunnett post-test with *p<0.05 and ***p<0.001. N = 2 (for basal) or 4 (for 3c) biological replicates. **(E)** Schematic of the approach taken to test for potential lasting effects of transient EHMT1/2 inhibition during reprogramming. **(F)** Quantification of viable pups obtained after blastocyst injection with indicated iPSCs. Each circle represents the offspring obtained from one recipient female of 20 injected blastocysts. ** = p<0.01 with Mann-Whitney test.

AA in recent years has been recognized as a potent modulator of the activity of chromatin-modifying enzymes (Cimmino et al., 2018; Monfort and Wutz, 2013), in particular of histone and DNA demethylases. In general, activation of AA-dependent enzymes is associated with successful nuclear reprogramming and iPSC formation (Chen et al., 2013b; Esteban et al., 2010; Tran et al., 2019). To explain our observation of impaired reprogramming in presence of AA, we speculated that EHMT1/2 might be required to balance the activities of specific enzymes that are stimulated by AA. In agreement with this notion, quantification of global H3K9me2 levels by immunofluorescence upon UNC0638 treatment revealed a slightly more pronounced reduction of this mark during enhanced than basal reprogramming (**Figure 1D**). We attempted to restore efficient reprogramming in presence of EHMT1/2 inhibitor by shRNA-mediated knockdown (KD) of the AA-stimulated H3K9 demethylases. Interference with KDM3B, which has recently been reported to be the main modulator of H3K9me2 downstream of AA in established pluripotent stem cells (Ebata et al., 2017), strongly impaired basal reprogramming but supported 3c enhanced reprogramming in absence of UNC0638 (**Figure 4C**). Knockdown of *Kdm3a* had no significant effects, despite similar reduction in mRNA abundance (**Figure S4C**). Importantly, interference with KDM3B also ameliorated the reduction in iPSC colony formation during 3c enhanced reprogramming in presence of UNC0638 (**Figure 4D**). These observations further suggest context-dependent functions of the H3K9 methylation machinery during iPSC formation and support the notion that the balance between EHMT1/2 and specific AA-stimulated enzymes is required for efficient reprogramming. Of note, while we have previously shown that 3c reprogramming yields developmentally highly competent iPSCs (Amlani et al., 2018), iPSCs derived from cells exposed to EHMT1/2 inhibition during the first 48 hours of reprogramming gave rise to significantly reduced numbers of viable pups upon blastocyst injections (**Figure 4E,F**). This raises the possibility that even transient interference with the balancing activity of these enzymes can have lasting adverse effects on the biological properties of iPSCs.

## DISCUSSION

A plethora of experimental evidence has demonstrated that H3K9 methyltransferases can counteract the induction of pluripotency ex vivo. By studying the consequences of EHMT1/2 inhibition in two well-defined reprogramming conditions, our work provides several lines of insight into how interference with the H3K9 methylation machinery affects cellular and molecular changes during iPSC formation.

First, our work confirms that counteracting H3K9 methylation facilitates reactivation of the *Pou5f1* locus (Chen et al., 2013b; Sridharan et al., 2013), which is consistent with the developmental role of EHMT2 (Feldman et al., 2006). Second, EHMT1/2 appear to be involved in the early downregulation of fibroblast-associated genes, indicating that H3K9 methyltransferases can contribute to the early silencing of the somatic program (Li et al., 2017). Why this effect is restricted to specific loci and is substantially more pronounced during enhanced reprogramming remains to be determined. It is possible that enhanced reprogramming is associated with the upregulation or stabilization of a protein co-factor that can recruit EHMT1/2 to target gene loci, as has been reported for MYC in cancer cells (Tu et al., 2018). Third, EHMT1/2 have a highly context-dependent function during the mesenchymal-to-epithelial transition necessary for reprogramming. In basal reprogramming, inhibition of EHMT1/2 facilitates upregulation of epithelial markers such as CDH1 and EpCAM, suggesting that H3K9 methylation contributes to the stable silencing of additional pluripotency-associated gene loci in somatic cells beyond *Pou5f1*. In striking contrast, EHMT1/2 inhibition strongly counteracts epithelization during enhanced reprogramming. This might be a consequence of the impaired silencing of fibroblast-associated gene loci discussed above or reflective of interference with other aspects of the molecular change that distinguish enhanced from basal reprogramming. It is noteworthy that EHMT1 and EHMT2 appear to have distinct roles during fibroblast reprogramming in our system. These enzymes predominantly function as a heterodimer in embryonic stem cells (Tachibana et al., 2005), while distinct functions of these methyltransferases have been reported in other cell types (Battisti et al., 2016).

The requirement for EHMT1/2 during enhanced reprogramming appears to be caused by AA. To our knowledge, none of the prior studies reporting facilitated reprogramming upon interference with EHMT1/2 utilized AA, explaining why this surprising interaction has been missed. In addition, most efforts to improve iPSC formation by interfering with H3K9 methylation were done at late stages during reprogramming or by using partially reprogrammed cells (Chen et al., 2013b; Sridharan et al., 2013; Tran et al., 2015). Of note, a genetic screen for epigenetic regulators conducted during human iPSC formation (which are routinely derived and cultured in presence of AA) reported reduced reprogramming efficiencies upon knockdown of EHMT1 (Onder et al., 2012). In order to explain the context-dependent consequences of EHMT1/2 inhibition, we propose that a balance between EHMT1/2 and AA-stimulated enzymes such as KDM3B is important for iPSC reprogramming. Of note, the impact of TET1 on the success of iPSC formation is also modulated by the presence of AA (Chen et al., 2013a), raising the possibility that DNA demethylases might functionally interact with EHMT1/2 during enhanced reprogramming.

Our data also suggest that even transient inhibition of H3K9me2 methyltransferase activity during reprogramming results in impaired iPSC function, suggesting persistent epigenetic aberrations. The precise molecular abnormalities caused by EHMT1/2 inhibition remain to be determined, but this observation represents a cautionary note for potential undesired consequences when targeting chromatin modifiers to facilitate iPSC formation. In conclusion, our results show that H3K9 methyltransferases can function as flexible regulators of cellular reprogramming that impact not only molecular change during the process, but also properties of resultant iPSCs. The concept of a balance between EHMT1/2 and AA-stimulated enzymes demonstrated here during iPSC formation might be helpful in developing new strategies targeting diseases driven by the dysregulation of chromatin-modifying enzymes.

## ACKNOWLEDGEMENTS

We thank Konrad Hochedlinger and current and past members of the Stadtfeld and Apostolou labs for helpful suggestions on this manuscript and during the course of this project. We are grateful to the Cytometry and Cell Sorting Laboratory and the Genomics Core at NYU Langone for expert help with our experiments. M.S. was supported by the National Institute of Health (1R01GM111852-01). L.E. is a New York Stem Cell Foundation Druckenmiller Fellow.

## AUTHOR CONTRIBUTIONS

S.V. and M.S. conceived the study and supervised data analysis. S.V. performed the initial characterization of the effect of EHMT1/2 inhibition on reprogramming, conducted most reprogramming experiments and optimized and conducted the isolation of reprogramming intermediates for RNA-seq, A.P. conducted bioinformatic analysis with support by Y.G. and A.T., JMV conducted reprogramming experiments with individual compounds, H.W. generated and validated knockdown constructs, E.S., L.E. and B.A. provided cell lines and other reagents, V.S. and J.A.S. assisted with RNA-seq, S.T. advised on the biology of histone methyltransferases, S.K. conducted blastocyst injections, E.A. was involved in experimental planning and supervised data analysis, M.S. conducted reprogramming, flow cytometry and immunofluorescence experiments and wrote the manuscript with input from all other authors.

## METHODS

### Mice

Derivation, handling, and genotyping of reprogrammable mice (JAX011001) with the *Oct4-GFP* allele were described previously (Stadtfeld et al., 2010). All animal experiments were in accordance with the guidelines of the NYU School of Medicine Institutional Animal Care and Use Committee.

### Cell culture

MEF cultures were established by trypsin digestion of midgestation (embryonic day (E) 13.5– E15.5) embryos and maintained in DMEM supplemented with 10% FBS, L-glutamine, penicillin/streptomycin, nonessential amino acids and β-mercaptoethanol. Reprogrammable MEFs were heterozygous for *Rosa26-rtTA* and for *Oct4-GFP* and either heterozygous for an inducible OKSM allele (Stadtfeld et al., 2010) or homozygous for an inducible OKS allele (Borkent et al., 2016). Established iPSCs were cultured on irradiated feeder cells in KO-DMEM (Invitrogen) supplemented with L-glutamine, penicillin/streptomycin, nonessential amino acids, β-mercaptoethanol, 1,000 U/ml LIF and 15% FBS (“ESC medium”). Reprogramming was carried out as previously described (Vidal et al., 2014). Briefly, 50-500 cells/cm^2^ were seeded on a layer of irradiated feeder cells in ESC medium in the presence of 1 μg/ml Dox and, if applicable, L-ascorbic acid (50 μg/ml), CHIR99021 (3 μM) and TGF-b RI Kinase Inhibitor II (250 nM) (“3c”). If applicable, UNC0638 (1 μM) was added. Media was changed every other day. Colonies were scored visually either after alkaline phosphatase staining or using the Oct4-GFP reporter allele.

### Immunofluorescence

Cells grown on 24-well plates were fixed with paraformaldehyde (4%), permeabilized with Triton X-100 (0.5%), and stained in blocking buffer (5% goat serum, 2 mg/ml fish skin gelatin and 0.2% Tween20 in PBS) with primary antibodies against H3K9me1 (ab9045; 1:200), H3K9me2 (ab1220; 1:200), H3K9me3 (ab8898; 1:200), H3K4me2 (ab7766; 1:200), H3K4me3 (ab8580; 1:200), EHMT1 (ab41969; 1:200) or EHMT2 (C6H3, 1:50) for one hour at room temperature, following by staining with appropriate AlexaFluor555-conjugated secondary antibodies (1:1000). After counterstaining nuclei with DAPI, cells were imaged using a Neo 5.5 cSMOS camera (Andor). Imaged quantification was conducted in NIS elements. Statistical analysis was conducted in Prism (GraphPad).

### Tetraploid blastocyst injections

Zygotes were isolated from BDF1 females as previously described (Stadtfeld et al., 2012) and cultured overnight until they reached the 2-cell stage. One hour after electro-fusion, 1-cell embryos were separated from embryos that had failed to fuse, cultured for another 2 days and then injected with iPSCs. Viable pups were defined as those that survived for at least three days following birth. Statistical analysis was conducted in Prism (GraphPad).

### shRNA mediated knockdown

97-mer oligonucleotides against specific target genes were designed using the splashRNA algorithm (Pelossof et al., 2017), PCR amplified using the primers miRE-Xho-fwd and miRE-EcoOligorev and cloned into the miRE plasmid backbones (Fellman et al. 2013). Viruses were produced using the packaging vectors psPax2, MD2G and Pasha/DCr8. Knockdown was confirmed using quantitative PCR using gene-specific primers. All oligonucleotides are listed in Supplemental Table 5.

### Flow cytometry

Reprogramming cultures were harvested by incubation with pre-warmed 0.25% trypsin/1mM EDTA solution for 5 minutes at 37°C. Single-cell suspensions were obtained by repetitive pipetting and transfer through a 40 μm cell strainer. Cells were incubated with eFluor 450-conjugated anti-THY1, biotin-conjugated anti-CDH1 and PE/Cy7-conjugated anti-EpCAM, followed by incubation with Streptavidin-APC (all eBiosciences) and data acquired on a FACS LSR2 (BD Biosciences), using DAPI to identify dead cells. Data analysis was conducted with FlowJo (Tree Star) and with Prism (GraphPad).

### RNA library preparations and analysis

Total RNA, extracted from cells with the RNeasy Plus Kit (Qiagen), with RIN values > 8 were subjected to Automated TruSeq stranded total RNA with RiboZero Gold library preparation (Illumina). Single-end 50 bp reads were generated with HiSeq2500. RNA-seq raw sequencing data were aligned to mouse genome version mm10 with the tophat algorithm (version 2.1.0)(Kim et al., 2013) and the use of «--b2-very-sensitive» parameter. Samtools (version 1.8) (Li et al., 2009b) was used for data filtering and file format conversion. Aligned reads were assigned to exons with the use of the HT-seq count (version 0.5.4p3) algorithm (Anders et al., 2015) and the following parameters «-m intersection-nonempty». Differentially expressed genes were identified with the use of DESeq R package (R.3.4.4) (Anders and Huber, 2010), excluding genes with RPKM<1. R was used for PCA analysis of all genes whose expression was above 1 RPKM in at least one condition and for k-mean clustering on Z-transformed normalized expression levels of DEGs between MEFs and early reprogramming intermediates. Gene ontology analysis was conducted using Gorilla (Eden et al., 2009). Raw sequencing data are submitted in GEO under accession number GSE130490.

---

**Figure S1.**
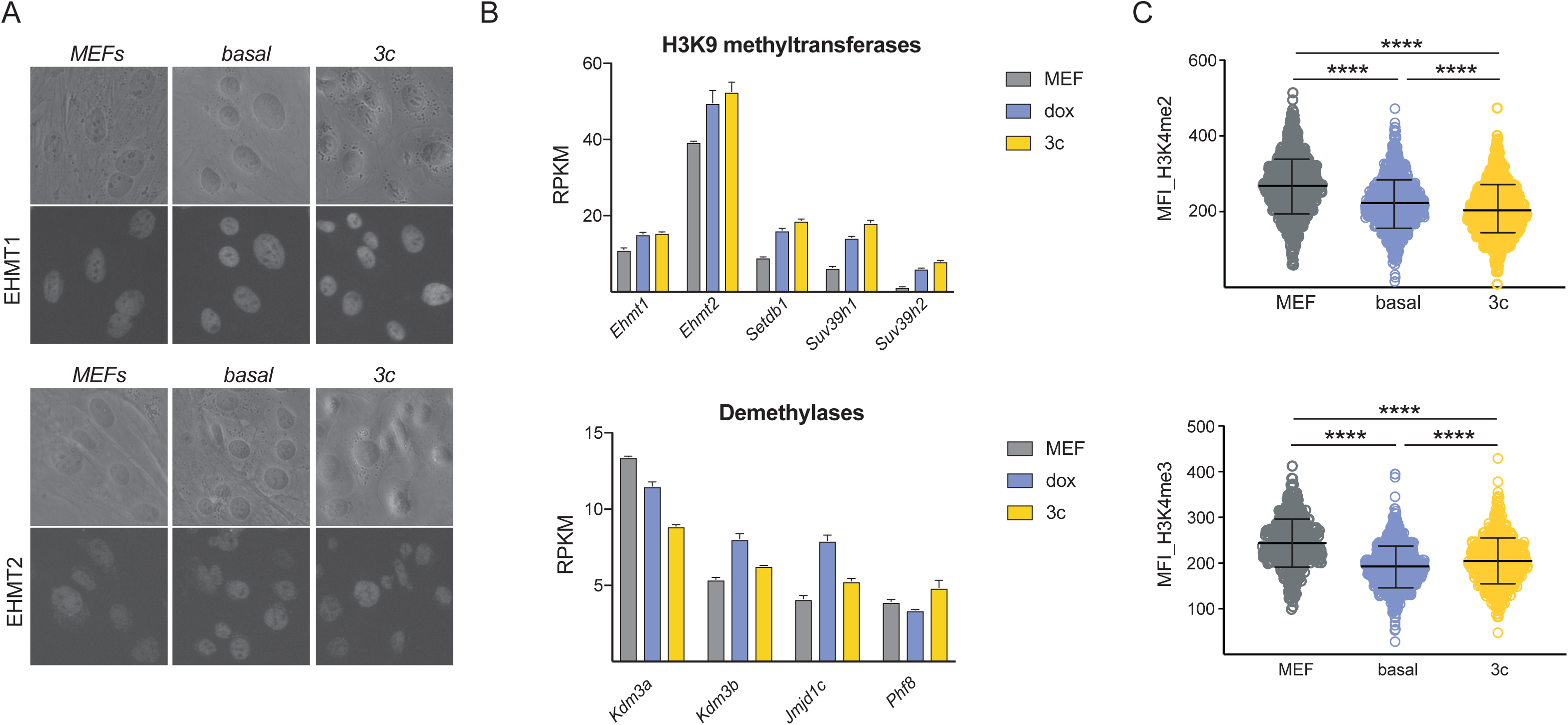
Increased EHMT1/2 activity during early stages of OKSM-driven reprogramming. **(A)** Representative brightfield and fluorescence images of MEFs and cells in indicated reprogramming conditions 24h after induction of OKSM and staining with antibodies against EHMT1 (upper) or EHMT2 (lower). **(B)** Expression levels of indicated H3K9 methyltransferases (upper panel) and demethylases (lower panel) as measured by RNA-seq in MEFs and during basal and 3c reprogramming (48h days of OKSM expression). **(C)** Quantification of H3K4me2 and H3K4me3 levels in indicated conditions. More than 200 nuclei of similar size were measured for each cell populations. Significance with one-way ANOVA with Tukey post-test with ****p<0.0001.

**Figure S2.**
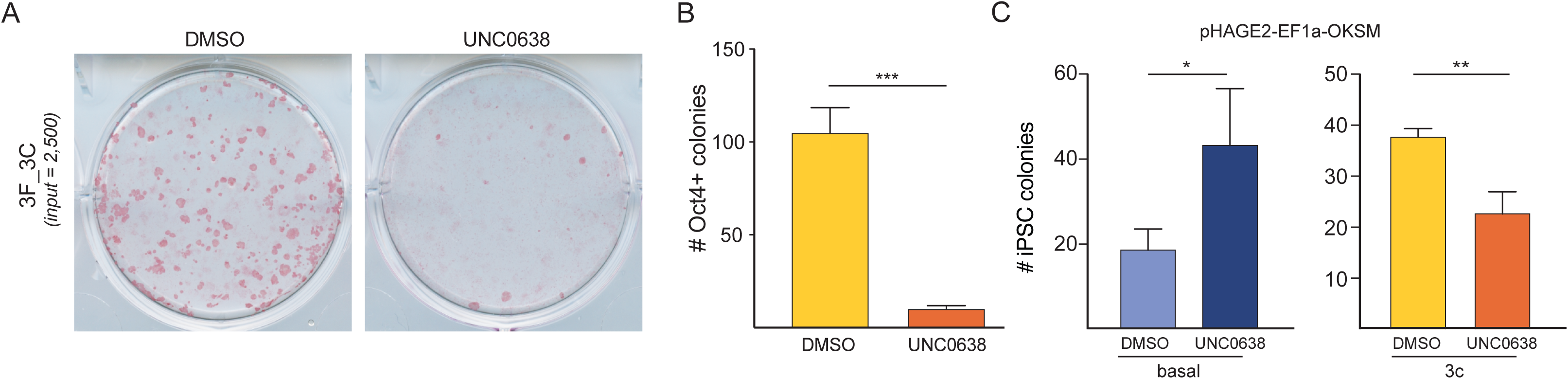
EHMT1/2 activity supports 3c enhanced reprogramming. **(A)** Representative AP staining of transgene-independent iPSC colonies obtained upon expression of OKS in MEFs for nine days in 3c enhanced reprogramming conditions in absence and presence of EHMT1/2 inhibition. **(B)** Quantification of iPSC colony formation upon OKS expression in indicate conditions. **(C)** Number of Oct4+ iPSC colonies formed upon transducing MEFs with a constitutive lentiviral vector expressing OKSM and reprogramming cells under basal or 3c enhanced conditions in absence of presence of EHMT1/2 inhibitor. Significance in (B-C) with two-tailed t-test with *p<0.05, **p<0.01 and ***p<0.001. N = 3 biological replicates.

**Figure S3.**
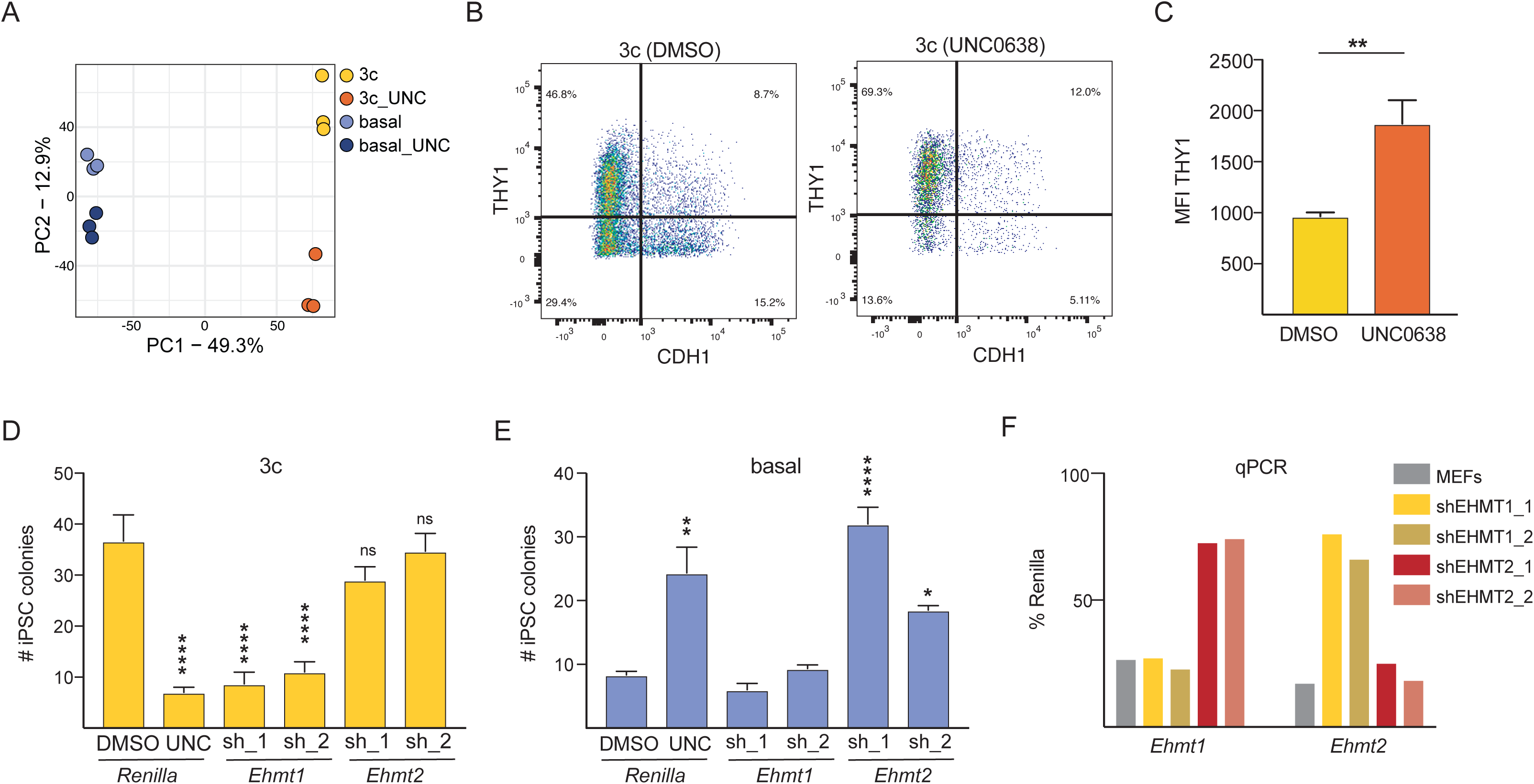
Molecular consequences of EHMT1/2 inhibition during early reprogramming stages. **(A)** Principal component analysis of RNA-seq data obtained from cells undergoing reprogramming for two days in indicated conditions. **(B)** Flow cytometry histograms showing expression levels of the fibroblast marker THY1 on CDH1+ cells at Day 4 of 3c enhanced reprogramming in absence and presence of EHMT1/2 inhibitor. **(C) (D)** Quantification of THY1 levels on CDH1+ intermediates at Day 4 of 3c enhanced reprogramming, MFI = mean fluorescence intensity (as measured by flow cytometry). Significance with t-test, **p<0.01. N = 3 biological replicates. **(D)** Number of stable iPSC colonies formed under 3c reprogramming conditions from cells transduced with indicated shRNAs. **(E)** Number of stable iPSC colonies formed under basal reprogramming conditions from cells transduced with indicated shRNAs. **(F)** mRNA levels of *Ehmt1* and *Ehmt2* in MEFs and in MEFs transduced with indicated shRNAs and undergoing reprogramming in 3c conditions for 48h relative to cells expressing shRNA targeting *Renilla*.

**Figure S4.**
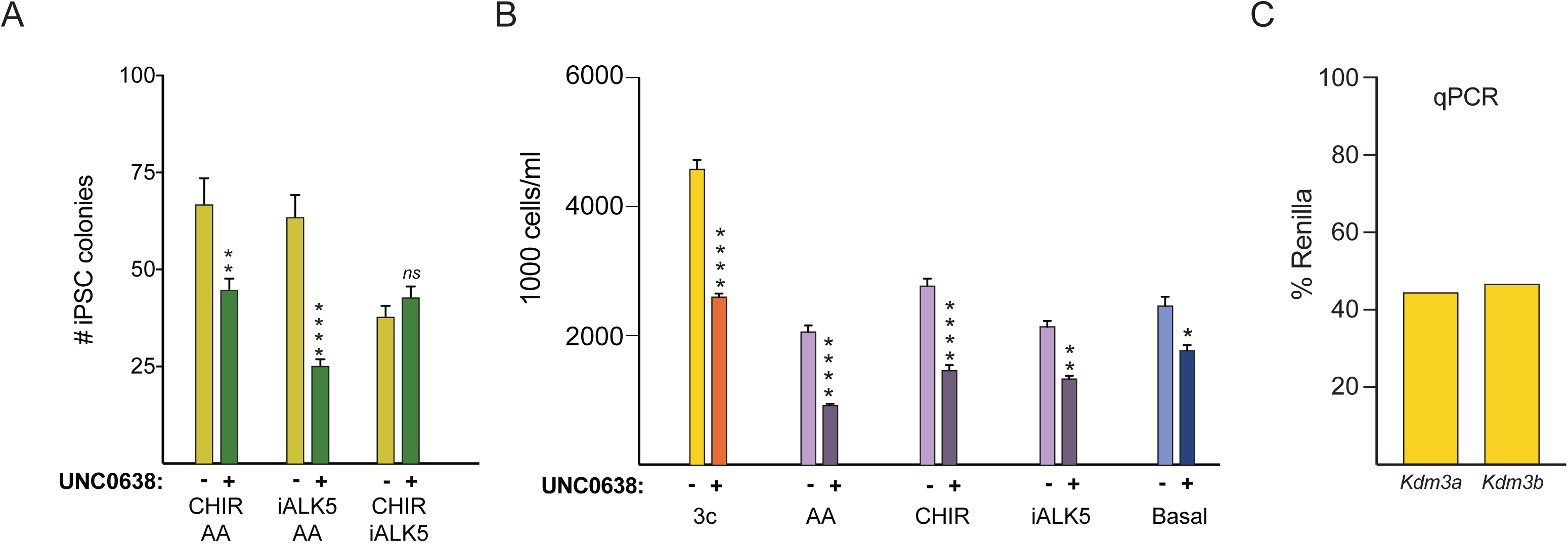
AA establishes a requirement for EHMT1/2 activity during enhanced iPSC reprogramming. **(A)** Quantification of iPSC colonies formed in presence of indicated dual compound conditions. N = 3 biological repeats. Significance was calculated by multiple t-test with p-values of **<0.01 and ****<0.0001. **(B)** Quantification of total cell numbers after expression of OKSM factors for 48h in indicated conditions in absence or presence of UNC. **(C)** mRNA levels of indicated histone demethylases in cells undergoing reprogramming in 3c conditions for 24h upon expression of respective shRNAs relative to cells expressing shRNA targeting *Renilla*.

